# Aging markers in human urine: A comprehensive, non-targeted LC-MS study

**DOI:** 10.1101/2020.05.10.086843

**Authors:** Takayuki Teruya, Haruhisa Goga, Mitsuhiro Yanagida

## Abstract

Metabolites in human biofluids document individual physiological status. We conducted comprehensive, non-targeted, non-invasive metabolomic analysis of urine from 27 healthy human subjects, comprising 13 youths (30±3 yr) and 14 seniors (76±4 yr). Quantitative analysis of 99 metabolites revealed 55 that were linked to aging, displaying significant differences in abundance between the two groups. These include 13 standard amino acids, 5 methylated, 4 acetylated, and 9 other amino acids, 6 nucleosides, nucleobases, and derivatives, 4 sugar derivatives, 5 sugar phosphates, 4 carnitines, 2 hydroxybutyrates, 1 choline, and 1 ethanolamine derivative, and glutathione disulfide. Abundances of 53 compounds decreased, while 2 increased in elderly people. Many age-linked markers were highly correlated; 42 of 55 compounds, showed Pearson’s correlation coefficients larger than 0.70. As metabolite profiles of urine and blood are quite different, age-related information in urine components offer yet more valuable insights into aging mechanisms of endocrine system and related organ systems.

## Introduction

Elderly people are acutely aware of the progressive aging of different body parts, but quantifying and characterizing physiological aging is less intuitive. Nonetheless, assessment of aging and determination of aging type by analyzing metabolites in biofluids, such as blood and urine, may help us to understand broadly-featured aging of the human body (Kampmann et al., 1974) (Ames, 1989) (Short et al., 2005) (Slupsky et al., 2007) (Lawton et al., 2008) (Mishur and Rea, 2012) (Yu et al., 2012) (Menni et al., 2013) (Gonzalez-Covarrubias et al., 2013) (Auro et al., 2014) (Chaleckis et al., 2016) (Hertel et al., 2016) (Jove et al., 2016) (Rist et al., 2017) (Chak et al., 2019). Age-dependent changes of metabolite abundances may be valuable to determine the molecular causes of impaired organ functions. Technology that enables simple and rapid measurement of urinary metabolites, which can be collected non-invasively, has certain advantages over methods using blood. Urinary metabolites are promising biological samples for monitoring health parameters if the metabolic processes resulting in production of those metabolites can be fully understood. To date, few comprehensive approaches to investigate human urinary metabolites in aging have been reported (Thevenot et al., 2015) (Hertel et al., 2016) (Rist et al., 2017). Urination is a primary route by which the body eliminates water-soluble waste products. Accordingly, urine has been broadly utilized for diagnosis of renal dysfunction in diverse kidney diseases (Han et al., 2002) (Eknoyan et al., 2003) (Mishra et al., 2005) (Parikh et al., 2006) (Pisitkun et al., 2006) (Vaidya et al., 2008). Nonetheless, because urinary metabolites originate in all organ systems, they may be useful to examine human aging. In this study, we analyzed urinary metabolites in elderly and young subjects, using comprehensive metabolomics to identify metabolites linked to aging. Striking correlations of many urine metabolites with age were found.

## Results

### Collection of urine samples

Samples of morning urine, immediately after awaking, were collected from healthy volunteer subjects (elderly, 75.8±3.9 yr, and young, 30.6±3.2 yr; **Supplementary Table s1** shows gender and BMI) in Onna Village, Okinawa, Japan. Precautions taken for sample collection are described in the Materials and Methods. Basic data analytical procedures were similar to those previously described (Chaleckis et al., 2016) (Teruya et al., 2019) (Kameda et al., 2020).

### 99 urine metabolites identified

Ninety-nine urinary metabolites, about half of which are amino acids and their derivatives, were identified and quantified using liquid chromatography – mass spectrometry (LC – MS) and MZmine 2 (Pluskal et al., 2010a) (Teruya et al., 2019) **(Supplemental Table s2).** These compounds were subdivided into 12 groups, containing 17 standard amino acids, 12 methylated amino acids, 6 acetylated amino acids and 15 other amino acids, 12 nucleosides, nucleobases, and derivatives, 4 sugar derivatives, 6 sugar phosphates, 3 vitamins and coenzymes (pantothenate, 4-aminobenzoate, nicotinamide), 4 choline and ethanolamine derivatives, 8 carnitines, 11 organic acids, and 1 antioxidant (oxidized form of glutathione, GSSG). Small amino acids, such as glycine and alanine, were not detected in our analysis due to the mass cutoff (100 m/z) used. Levels of individual compounds, categorized by abundance as H (high, >10^8^), M (medium, 10^7^∼10^8^) or L (low, <10^7^), were estimated based upon mass spectroscopic peak area (Chaleckis et al., 2014) (Chaleckis et al., 2016). Some compounds varied widely from one individual to the next and are denoted as H-L, H-M or M-L. According to abundance, there were 7 H, 19 H-M, 7 H-L, 5 M, 45 M-L and 16 L compounds. Twenty-six urinary metabolites were abundant (H or H-M), the great majority of which were amino acids and their derivatives, such as the methylated amino acids, betaine and dimethyl-arginine, but they also included nucleosides, such as pseudouridine and N-methyl-guanosine. In blood, sugar phosphates and methylated amino acids were enriched in red blood cells (Chaleckis et al., 2014) (Chaleckis et al., 2016). In urine, sugar phosphates are age-related, except for glucose-6-phosphate. We show below that about half of all urinary metabolites are age-related.

### Most urinary age markers decreased with age

Of 99 urinary metabolites assayed in 27 subjects, 55 showed statistically significant differences between young and elderly (p-values, 0.00003 < p < 0.05) (**Fig. 1**). Thus, about half of these compounds are affected by aging, with most becoming less abundant in elderly subjects. Two exceptions, myo-inositol and glutathione disulfide (GSSG), were more abundant in elderly samples. In other words, ∼50% of all urinary metabolites decline in old age. Five of 55 metabolites, creatinine, dimethyl guanosine, decanoyl-carnitine, N-acetyl-aspartate, and tryptophan, were previously reported as urinary age markers (Kampmann et al., 1974) (Slupsky et al., 2007) (Thevenot et al., 2015) (Rist et al., 2017). myo-Inositol and GSSG increased 1.34- and 6.96-fold, respectively (**Table 1**). The increase of GSSG was striking, though its abundance in urine was rather low. Many metabolites diminished to less than half in urine of elderly subjects [fructose-1,6-diphosphate (0.14), carnosine (0.34), glycerophosphate (0.32), 2- and 3-hydroxybutyrate (0.33, 0.27, respectively), and octanoryl-carnitine (0.40)].

**Table 1.** List of 55 aging markers.

| *Category /Compound* | Peak<br>abundance | Ratio<br>(elderly/young) |
| --- | --- | --- |
| ***Nucleosides, nucleobases, and derivatives (6)*** |  |  |
| Dimethyl-guanosine | H-M | 0.63 |
| N-Methyl-adenosine | H-M | 0.56 |
| Pseudouridine | H-M | 0.45 |
| N-Methyl-guanosine | M | 0.51 |
| Adenosine | M-L | 0.61 |
| Hypoxanthine | M-L | 0.50 |
| ***Sugar derivatives (4)*** |  |  |
| Gluconate | H-M | 0.67 |
| N-Acetyl-glucosamine | H-M | 0.70 |
| 1,5-Anhydroglucitol | M-L | 0.45 |
| myo-Inositol | M-L | 1.34 |
| ***Sugar phosphates (5)*** |  |  |
| Glycerol-phosphate | H-L | 0.32 |
| Fructose-1,6-diphosphate | L | 0.14 |
| Fructose-6-phosphate | L | 0.33 |
| Pentose-phosphate | L | 0.65 |
| Phosphoglycerate | L | 0.46 |
| ***Choline and ethanolamine derivatives (2)*** |  |  |
| Glycerophosphocholine | M-L | 0.50 |
| Glycerophosphoethanolamine | L | 0.44 |
| ***Carnitines (4)*** |  |  |
| (iso)Butyryl-carnitine | H-M | 0.61 |
| (iso)Valeryl-carnitine | H-L | 0.66 |
| Decanoyl-carnitine | M-L | 0.49 |
| Octanoyl-carnitine | M-L | 0.40 |
| ***Organic acids (2)*** |  |  |
| 2-Hydroxybutyrate | M-L | 0.33 |
| 3-Hydroxybutyrate | M-L | 0.27 |
| ***Antioxidant (1)*** |  |  |
| Glutathione disulfide | L | 6.96 |
| ***Standard amino acids (13)*** |  |  |
| Histidine | H-M | 0.68 |
| Phenylalanine | H-M | 0.53 |
| Tryptophan | H-M | 0.49 |
| Asparagine | M-L | 0.73 |
| Glutamine | M-L | 0.46 |
| Leucine | M-L | 0.42 |
| Lysine | M-L | 0.50 |
| Methionine | M-L | 0.74 |
| Serine | M-L | 0.54 |
| Threonine | M-L | 0.71 |
| Aspartate | L | 0.62 |
| Isoleucine | L | 0.38 |
| Valine | L | 0.30 |
| ***Methylated amino acids (5)*** |  |  |
| Dimethyl-arginine | H | 0.69 |
| N-Methyl-histidine | H | 0.64 |
| Trimethyl-lysine | H | 0.74 |
| Dimethyl-lysine | H-M | 0.52 |
| Butyro-betaine | M-L | 0.76 |

**Table 1. List of 55 aging markers. (continued)**
| *Category* /Compound | Peak<br>abundance | Ratio<br>(elderly/young) |
| --- | --- | --- |
| **Acetylated amino acids (4)** |  |  |
| N-Acetyl-arginine | H-M | 0.78 |
| N2-Acetyl-lysine | M | 0.62 |
| N6-Acetyl-lysine | M | 0.51 |
| N-Acetyl-aspartate | M-L | 0.50 |
| **Other amino acids (9)** |  |  |
| Creatinine | H | 0.43 |
| Indoxyl-sulfate | H | 0.56 |
| 4-Guanidinobutanoate | H-M | 0.46 |
| Taurine | H-M | 0.52 |
| Carnosine | M-L | 0.34 |
| Keto(iso)leucine | M-L | 0.36 |
| Kynurenine | M-L | 0.45 |
| Quinolinic acid | M-L | 0.44 |
| S-Adenosyl-homocysteine | L | 0.75 |

The list of 55 aging-linked compounds that showed a significant difference in abundance between young and elderly people. The abundance of compounds (peak area) displayed: H, high peak areas [ $>10^8$ AU (arbitrary unit)]; M, medium peak areas ( $10^8 \sim 10^7$ AU); L, low peak areas ( $<10^7$ AU). The peak ratio was calculated using the median of peak abundance in elderly and young people, respectively.

**Figure 1.**
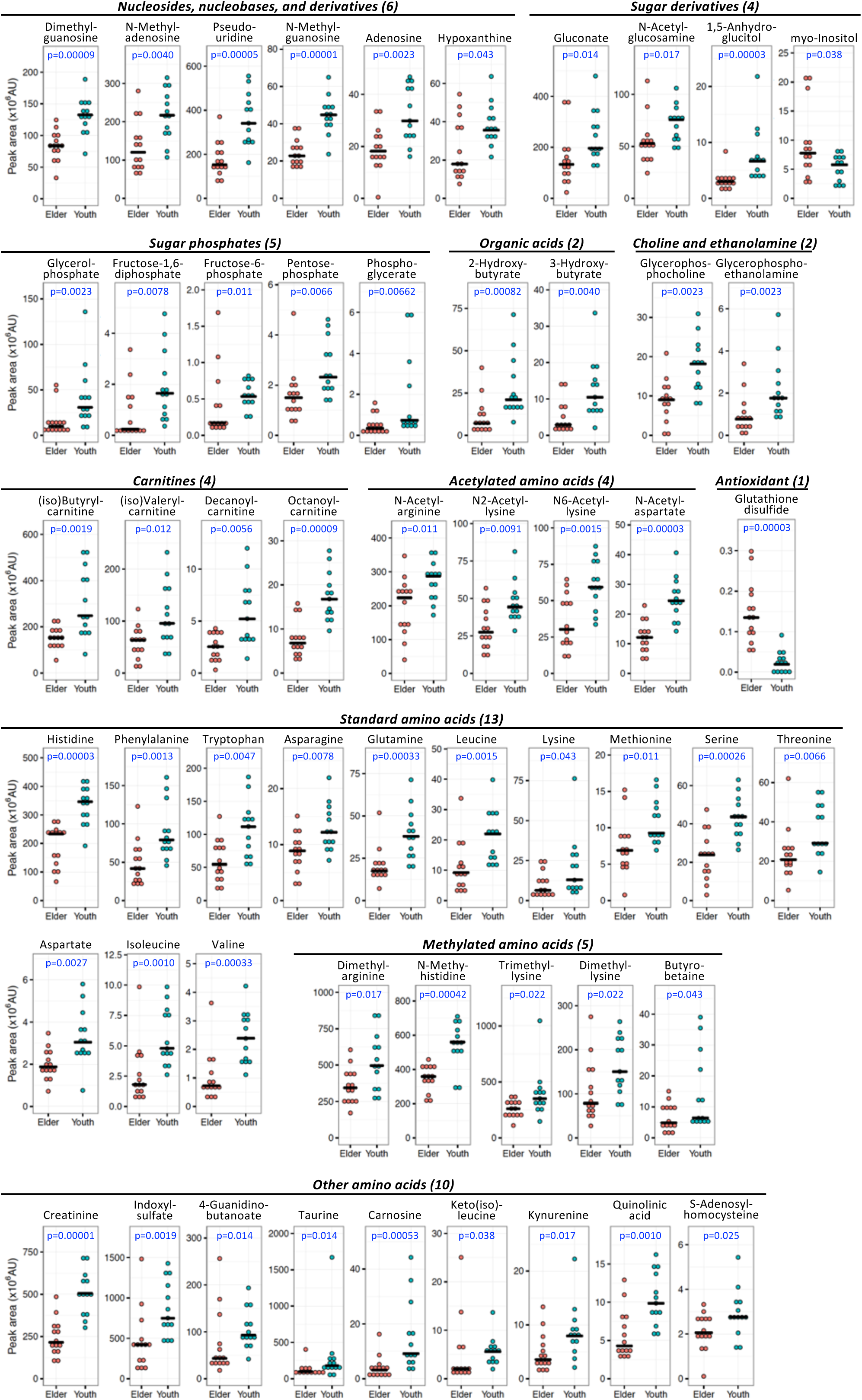
Dot plots of 55 compounds that showed significant differences in abundance between elderly and young people. Distributions of 11 subgroups of 55 metabolites in urine samples from 27 individuals. Pale red and azure dots represent elderly and young subjects, respectively. Bars represent medians in each group. The ratio of median value between elderly and young is shown in **Table 1.** P-values were obtained using the non-parametric Mann Whitney U test. Fifty-three of 55 compounds were more abundant in young subjects, while two (myo-inositol and GSSG) were more abundant among the elderly.

### Highly correlated age-linked urinary metabolites

To understand relationships among urine metabolites, Pearson’s correlation coefficients were calculated from abundance data for all 55 age-linked urinary metabolites of all 27 subjects (Methods section). The highest correlation, 0.92, was obtained for isoleucine – leucine and N-methyl guanosine – dimethyl guanosine (**Fig. 2A**). This is probably due to their similar chemical structures and proximity in biochemical pathways (see below and KEGG, https://www.genome.jp/kegg/). However, correlation values 0.91 obtained for glycerol-phosphate – 3-hydroxybutyrate, pseudouridine – 2-hydroxybutyrate, and pseudouridine – isoleucine clearly have a different explanation. Fifteen urinary metabolites having correlation values >0.85 formed a network (**Fig. 2B**). The high correlation between pseudouridine and isoleucine seems to be a key connection between two groups of metabolites. Additionally, the connections of creatinine to pseudouridine and 3-hydroxybutyrate to glutamine appeared to be required for further correlation (see below). In the case of a creatinine-related network, 15 metabolites have correlation coefficients from 0.71∼0.84 (**Fig. 2C**). Creatinine, a known waste product from muscle (Heymsfield et al., 1983) (Baxmann et al., 2008), is correlated with many metabolites. It even shows a negative correlation with GSSG (**Fig. 2D**).

**Figure 2.**
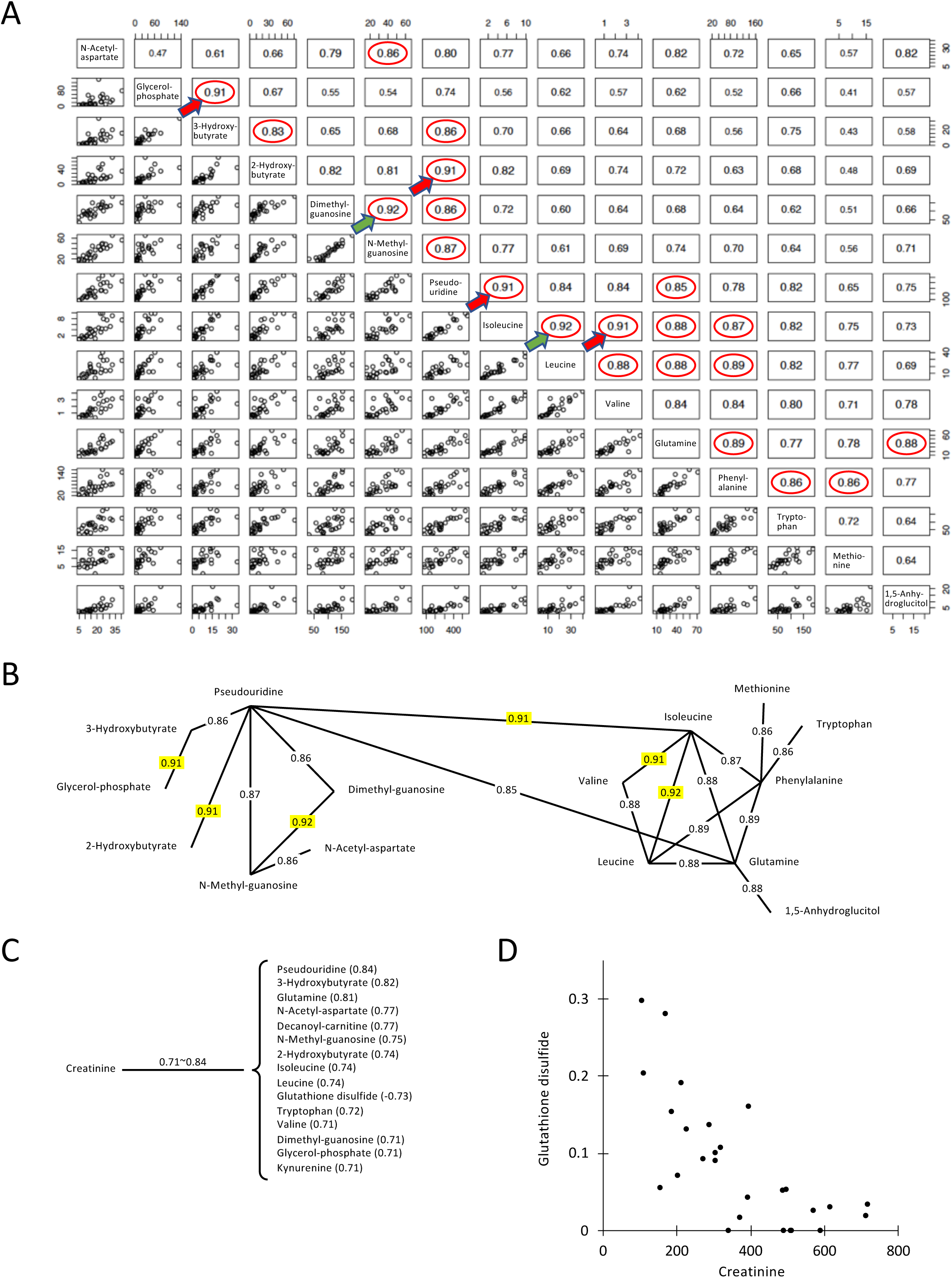
Correlation network among age-linked compounds. (A) Values indicate correlation coefficients between paired compounds. Highly correlated pairs of aging markers (r>0.85) are indicated with red circles. The most highly correlated pairs are indicated by green (0.92) and red arrows (0.91). (B) Interrelated compounds form a network. Correlations (r>0.90) are highlighted in yellow. (C) Fifteen metabolites with the highest correlation with creatinine are listed. (D) Scatter plot of the peak abundance (x10^6^ AU) between creatinine and GSSG. GSSG was negatively correlated with creatinine.

Pairwise correlation analyses revealed 21 pairs of compounds with correlation coefficients >0.85 (**Fig. 2A**). These show a network consisting of two groups, one centered around pseudouridine and the other around isoleucine (**Fig. 2B**). This suggests that compounds within each of these groups may be metabolically linked.

When urinary compounds with correlation coefficients larger than 0.7 were selected, the majority of age-linked metabolites formed a large network consisting of 42 metabolites (**Supplemental Fig. S1)**. Since all age-linked metabolites displayed p-values < 0.05 in dot plot profiling, the fact that the great majority (42/55 = 76%) form such a large age-linked correlation network is quite impressive.

### Some age-linked metabolites were not highly correlated

On the other hand, 13 compounds, though they were age-related, showed only weak correlations (<0.7) with the other 42 age-linked urinary metabolites (**Supplemental Fig. S1**). Aspartate, a standard amino acid having an acidic side chain, showed maximal, but inverse correlation with GSSG (−0.67). Aspartate has a correlation coefficient of 0.65 with N-acetyl aspartate, indicating that structural similarity might partly explain the weak correlation. S-adenosyl-homocysteine did not show correlation values higher than 0.41 (N-methyl-adenosine). Although the correlation is rather low, the two compounds exhibit structural similarity. Similarly, lysine, dimethyl-lysine, trimethyl-lysine, and N6-acetyl-lysine did not show the correlation coefficients higher than 0.5 among them. These metabolites diminish significantly in urine of elderly people. Fructose-1,6-diphosphate showed a maximal correlation value (0.58) with pentose-phosphate and gluconate, all carbohydrates, but correlations were low (**Supplemental Fig. S2**). Whether these metabolites decline in elderly subjects as a consequence of aging or as a cause of it, remains to be investigated.

### Heatmap analysis of urinary metabolites linked to aging

We then employed a heatmap to visualize the quantitative profile of age-linked urinary metabolites in individual subjects (**Fig. 3**). Since most of these metabolites declined in the elderly, matrix color represents metabolite abundance for individual subjects (t-score<40 white, low; 40∼50 thin red, slight low; 50∼60 moderate red, slight high; >60 deep red, high). Elderly samples to the left were mostly white or pale red (lower level), except for myo-inositol and GSSG, which were moderate or deep red (higher level). Thus, a heatmap of urinary metabolites graphically illustrates the degree of urinary metabolite aging for individual subjects, so that subjects may be compared one with another. Two subjects (elderly 11, 81F and young 15, 33M) showed patterns remarkably like those of the opposite age group.

**Figure 3.**
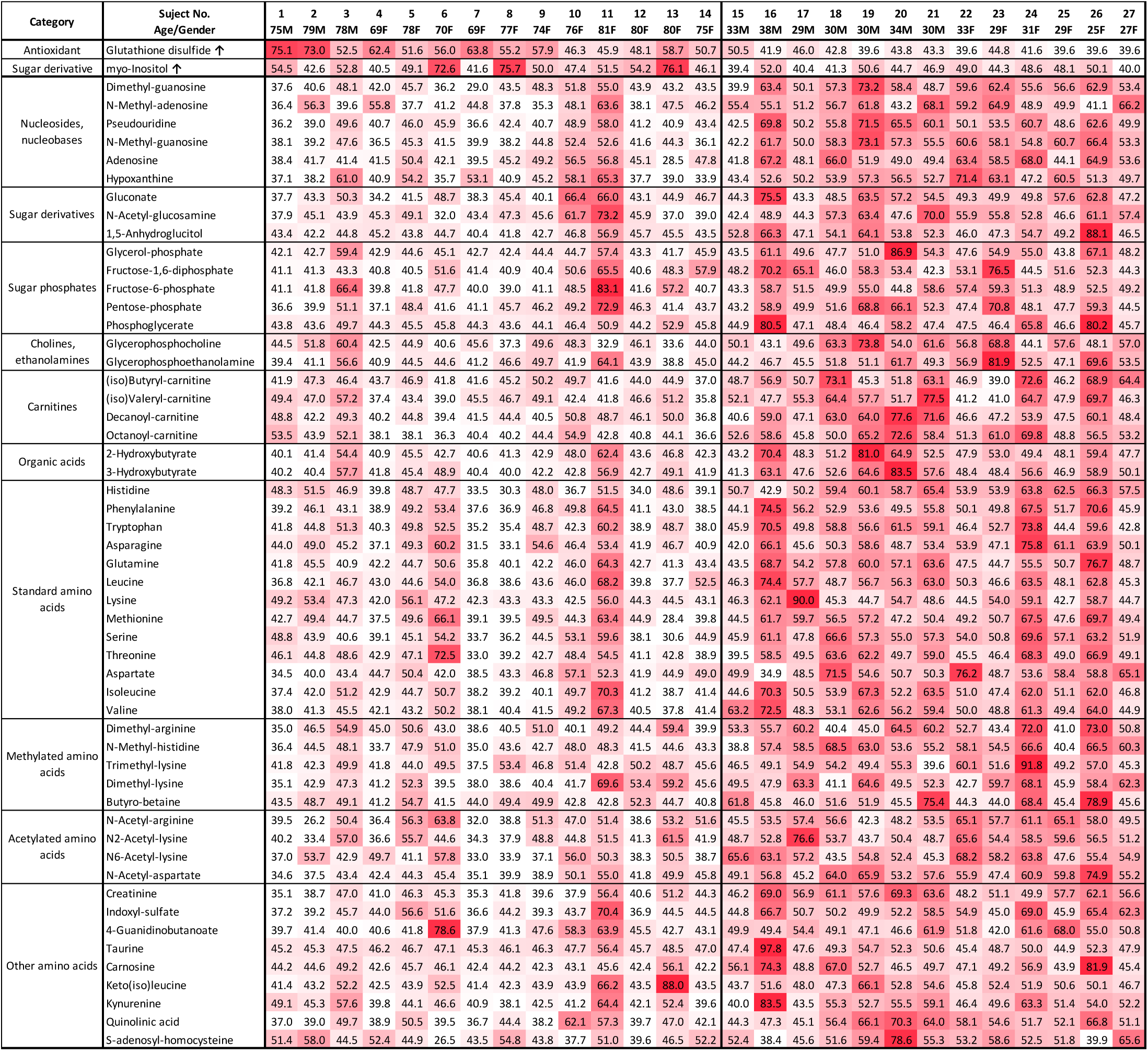
Heatmap representing urinary metabolic profiles of elderly and young subjects. Standardized data for each metabolite are shown for 27 subjects using a color matrix representing relative abundance data of 55 urinary aging markers. Numerical values indicate the t-score, a kind of standardized score. The mean and standard deviation are 50 and 10, respectively. Color intensity of the cells reflects the t-score, indicating levels higher than average.

### Principal component analysis of age-linked metabolites

Implications of urinary compounds in aging should also be cross examined using results of blood compound analyses. However, as these compounds decrease in urine of elderly people, the results are consistent with actual aging. Some urinary metabolites may be implicated in sustaining health and slowing aging. To integrate quantitative metabolite abundance data from individual subjects, abundances of these 55 age-related metabolites were subjected to principal component analysis (PCA) (Nakamura et al., 1988) (Kameda et al., 2020). Subjects were separated into 2 groups (red for elderly and blue for young subjects) represented by negative and positive values of principal component (PC) 1, respectively (**Fig. 4A**). One elderly subject 11 (81F) was in the middle of the young group and one young subject 15 (33M) was in the group of elderly subjects in the PCA plot (**Fig. 4A**). These are consistent with results plotted in the heatmap (**Fig. 3**). The PC1 score and chronological age of the subject showed a high correlation with r=0.80. (**Fig. 4B**).

**Figure 4.**
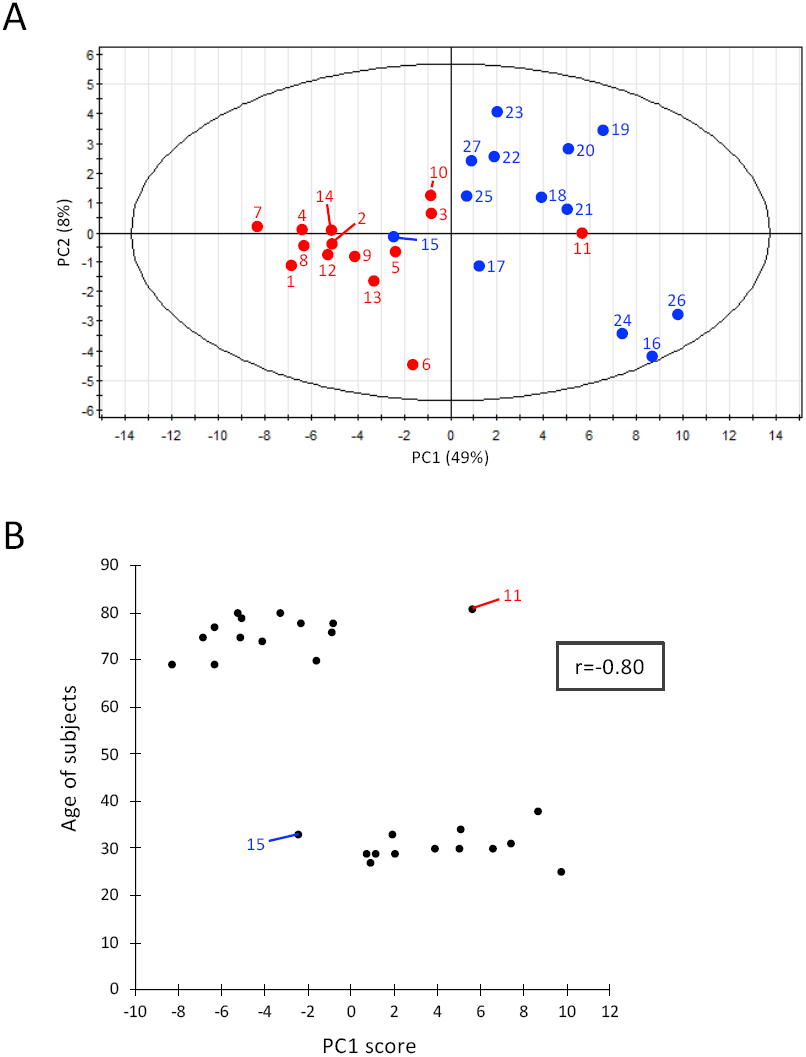
PCA of 55 aging markers. (A) Thirteen young and 14 elderly subjects are shown in blue and red, respectively, with their subject numbers (corresponding to **Fig. 3**). PC2 comprised metabolites that were not strongly correlated (glycerophosphocholine, S-adenosyl-homocysteine, etc.), but that were isolated from a strong correlation network. (B) PC1 score of each subject (X-axis) is plotted versus subject age (Y-axis). The correlation coefficient is shown in the box.

### Negative correlation between GSSG and age-linked metabolites

GSSG showed significant negative correlations with some age-linked metabolites (N-acetyl aspartate, N-methyl guanosine, dimethyl guanosine, pseudouridine, quinolinic acid, creatinine, N-methyl-histidine, etc.) (**Fig. 5A**). While the oxidized disulfide form was detected in elderly samples, but the reduced form was not detected in young or elderly samples. Thus only the inactive form of glutathione was detected in elderly samples. A possible interpretation is that the active reduced form glutathione (GSH) may be very reactive and short-lived as a metabolite (see Discussion). The abundance of GSSG in 27 subjects is shown together with N-acetyl aspartate, N-methyl guanosine, dimethyl guanosine, pseudouridine, and chronological age of subject (**Fig. 5B-F**). GSSG was low to undetectable in 5 young subjects, but accumulated in the elderly samples (**Fig. 5F**).

**Figure 5.**
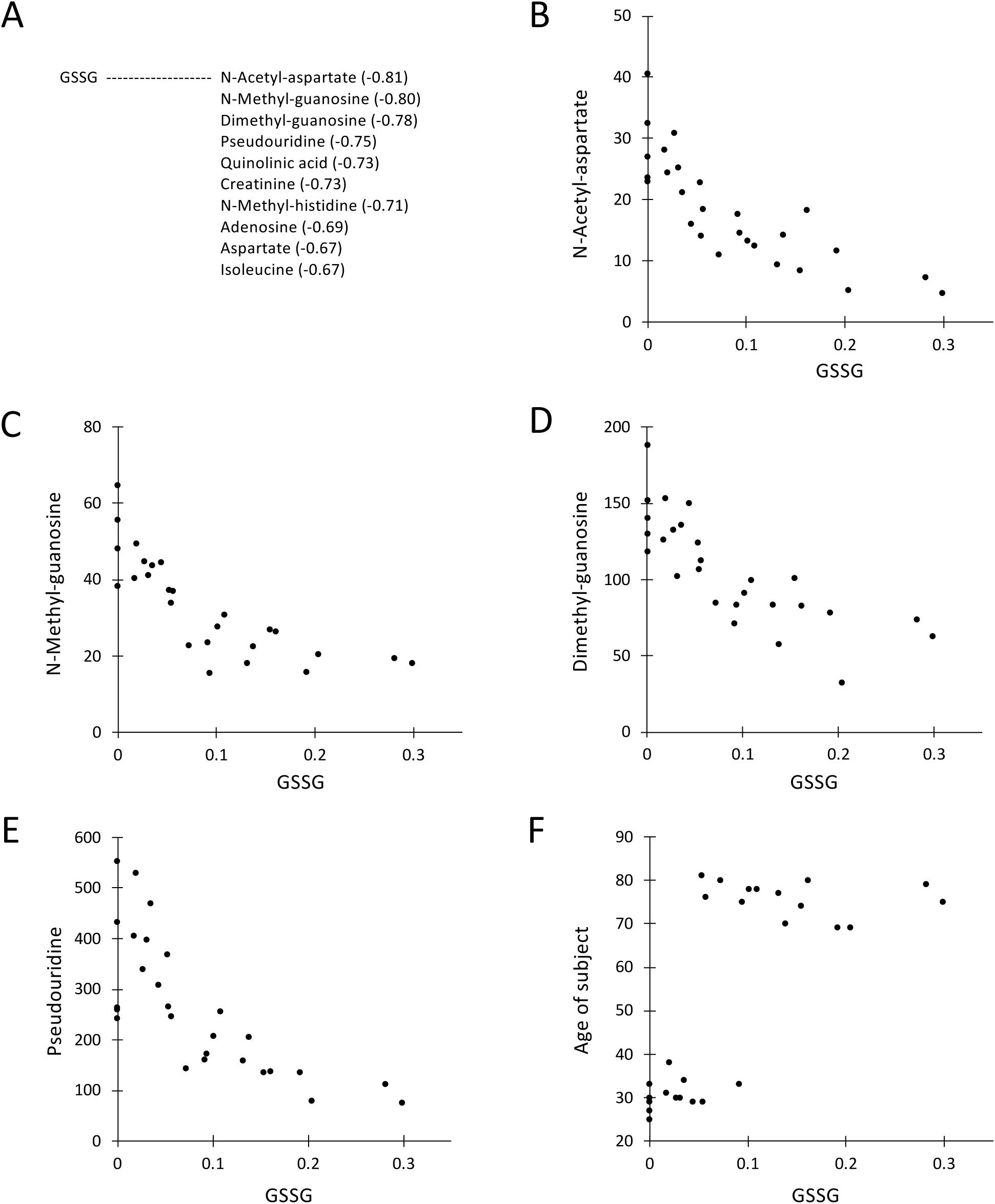
Scatter plots between GSSG and correlated compounds or age. (A) Ten metabolites with the highest correlations with GSSG are listed. Scatter plot of the peak abundance (x10^6^ AU) between GSSG and N-acetyl aspartate (B), N-methyl guanosine (C), dimethyl guanosine (D), pseudouridine (E), and the age of the subject (F).

GSSG showed different degrees of abundance in different subjects, and was virtually absent in young subjects, but in elderly subjects, its abundance increased and paralleled the abundance of creatinine in young subjects. Thus GSSG abundance was inversely related to aging. The reason for this increase of GSSG in elderly urine is probably due to the absence or declining effectiveness of a mechanism to metabolize and reprocess GSSG by the elderly. Such a reductive mechanism does exist in young subjects; however, the redox environment regarding glutathione appears to be altered during aging. Inability to reduce oxidized GSSG may accelerate aging.

## Discussion

We initially investigated metabolomics of a simple model eukaryote, fission yeast (*Schizosaccharomyces pombe*) using yeast genetic technology to identify and determine metabolite profiles in wild type and mutant cell extracts (Pluskal et al., 2010b) (Pluskal et al., 2016) (Pluskal and Yanagida, 2016a) (Pluskal and Yanagida, 2016b). By adapting the software MZmine 2 for metabolite identification (Pluskal et al., 2010a) (Pluskal et al., 2012), we found that metabolomics of fission yeast and humans are surprisingly similar in regard to metabolite composition (Chaleckis et al., 2014). Metabolites detected in fission yeast and human whole blood were 75% identical. Given this unexpected similarity, we adapted our techniques to human blood metabolites to better understand health, disease, and longevity (Chaleckis et al., 2014) (Chaleckis et al., 2016) (Teruya et al., 2019) (Kameda et al., 2020).

### Distinction of aging information between blood and urine

Urinary metabolites offer a non-invasive means of obtaining aging information. The present results indicate that urinary aging information may be distinct from that obtainable from blood. Blood metabolites derived from plasma and red blood cells of elderly subjects reflect a decrease in antioxidant production and muscle activity or increasing inefficiency of nitrogen metabolism (Chaleckis et al., 2016). Aging information obtained from urine will be useful, as human aging is exceedingly complex. In the present study, we identified 55 human urinary aging markers using non-targeted, comprehensive LC-MS. Since urine contains 99 compounds, many urinary metabolites such as vitamins (pantothenate and nicotinamide) and organic acids (citrate, malate) neither significantly decreased nor increased so that about a half of urinary metabolites are not age markers. In aged subjects, levels of protein, nucleic acid and lipid synthesis, modification, and turnover may decrease due to reduced physical activity (Hughes et al., 2002) (Pollack et al., 2002) (Maynard et al., 2015), resulting in declining metabolite abundances.

Metabolite data may be considered as an overview of the physiological state of all tissues and organs. Hence these 55 metabolites constitute a panorama of human aging as seen through urine composition. Alternatively, these may not represent actual human aging, but may represent voluntary lifestyle changes that may be reversible regardless of actual age.

### Myoinositol, GSSG and pseudouridine

Two exceptional urinary age metabolites, myo-inositol and presumably inactive, oxidized GSSG increased in elderly samples (1.34- and 6.96-fold, respectively). Myo-inositol becomes an important second messenger if phosphorylated, causing changes in [Ca^2+^] (Berridge, 1993) (Berridge, 2016). In the correlation analysis, myo-inositol was not highly correlated with any other metabolite. Glycerophosphocholine and N-methyl-adenosine showed weak, negative correlations with myo-inositol (−0.47and -0.43, respectively, **Supplemental Fig. S2**). Implications of the myo-inositol increase in elderly urine remain unclear. GSSG that we detected was the inactive, oxidized form. It increased in elderly urine. The active SH form of glutathione seems to be very unstable and it is difficult to prevent its degradation during preparation of urine as well as blood samples (Chaleckis et al., 2016, the present study). The active SH form was rapidly oxidized to an inactive disulfide form, which accumulated in elderly subjects, whereas in young subjects the disulfide form was hardly formed or undetected possibly due to rapid decay in the young body or sample. Consistently, reduced GSSG was undetectable in urine of the five young subjects **Fig. 2D** and **Fig. 5B-F**). The abundance of GSSG was inversely related to those of pseudouridine, creatinine, and 2-hydroxybutyrate, which were most abundant in young subjects. Thus GSSG in urine appears to be appropriate as an age marker in urine of elderly subjects. In contrast, the high abundance of pseudouridine seems to be emblematic of youth.

### Highly correlated urine metabolites

Fifteen metabolites are highly correlated (correlation coefficients >0.85, **Fig. 2A, B**). They consisted of two subgroups. One comprised nucleosides pseudouridine, dimethyl guanosine, N-methyl guanosine and 2- or 3-hydroxybutyrate and the other group contained branched-chain amino acids and aromatic amino acids. Correlation within each group was high, but was not always high between the two subgroups (0.65 for pseudouridine and methionine, and 0.56 for glycerol phosphate and isoleucine; **Supplemental Fig. S2**). It remains to be determined what kind of functional distinctions exist between these two subgroups of metabolites in health and disease. In addition, a number of compounds showing correlation coefficients >0.70 are structurally only remotely related. Therefore, the high correlations of urinary compounds cannot all be easily explained. While isoleucine, leucine, and valine contain hydrophobic side chains, glutamine, having a hydrophilic sidechain, showed correlation coefficients of 0.88-0.89 with isoleucine. Pseudouridine was highly correlated with 10 other compounds (>0.82), although we are unable to offer an explanation for this at present. The high correlation (0.83) between 2-hydroxybutyrate and 3-hydroxybutyrate may be explained by their structural relatedness, but their correlations in the range of 0.6-0.82 with 10 other compounds are difficult to explain.

### Thirteen metabolites linked by low correlation

Thirteen metabolites linked by low correlation coefficients are shown in **Supplemental Fig. S2.** They include myo-inositol, two regular amino acids (aspartate and lysine), methylated or acetylated amino acids, etc. Creatinine, an abundant aging marker (Kampmann et al., 1974) and an amino acid waste product in muscle (or other tissues) showed moderate correlations (**Fig. 2C**). Its urinary content decreased to 43% in elderly samples (**Table 1**). More than half (31) of the declining age markers in urine were amino acids (standard, methylated, acetylated, and other), and 6 more are nucleosides, nucleobases, and their derivatives. Protein degradation and nucleic acid turnover seem to cause the change in elderly urine. Pseudouridine is a tRNA component (Charette and Gray, 2000), perhaps important in catabolism of nucleosides and other compounds, as it is highly correlated with purine nucleosides (adenosine, guanosine, and inosine), muscle amino acids (isoleucine, creatinine), and organic acids (hydroxybutyrates, glycerol-phosphate). Pseudouridine is also abundant in blood, but the amount in urine is 15-fold higher. Pseudouridine showed very high correlation coefficients with 10 compounds (**Fig. 2A, B**) and its high abundance in urine may provide a convenient youth marker.

### An effective overview on aging by heatmap

We showed that heatmap analysis of age-linked metabolites provides an effective overview of aging patterns using urinary metabolite profiles of individual subjects. Heatmap patterns are convenient for visualizing differences between young and aged subjects and also individual variations within and between groups (**Fig. 3**). PCA was useful to categorize a group of subjects because of its capacity to integrate a large collection of data about various metabolites. Elderly and young subjects were clearly separated into two populations, with two exceptions (**Fig. 4**). Exceptional individuals are of considerable interest, regarding their persistent youth or premature aging, if these truly represent metabolic features. Their individuality may be worthy of further investigation. Among 55 age-related urinary markers, 13 compounds did not show correlations >0.7 (**Supplemental Fig. S1**). We determined the correlations of these 13 compounds with all metabolites. Age-related metabolites such as myo-inositol, S-adenosyl-homocysteine, N-methyl-adenosine, lysine, trimethyl-lysine, fructose-1,6-diphosphate, and glycerophosphocholine are only weakly correlated (∼0.6) (**Supplemental Fig. S2**). Thus, high correlation is not necessarily required of age-related compounds. These metabolites that are only remotely correlated are of interest to better understand the metabolic breadth of human aging.

## Materials and Methods

### Participants and sample collection

14 elderly (69∼81 yr) and 13 young (25∼38 yr) healthy people participated as subjects in this study (**Supplemental Table S1**). Measurements of metabolites in first morning urine are more consistent than random daytime sampling to monitor metabolites. Intra-individual coefficients of variation in first morning urine and 24-h collected urine are reportedly similar (Witte et al., 2009).

After collection, urine samples were brought to the laboratory within 3 hr 0.2 mL urine were immediately quenched in 1.8 mL of 55% methanol at -40°C. This quenching step stabilizes metabolites and maximizes reproducibility of metabolomic data. Two internal standards (10 nmol of HEPES and PIPES) were added to each sample. After brief vortexing, samples were transferred to Amicon Ultra 10-kDa cut-off filters (Millipore, Billerica, MA, USA) to remove proteins and cellular debris. After sample concentration by vacuum evaporation, each sample was re-suspended in 40 μL of 50% acetonitrile, and 1 μL was used for each injection into the LC-MS system, as described (Chaleckis et al., 2016) (Kameda et al., 2020).

### Ethics statement

Written, informed consent was obtained from all donors, in accordance with the Declaration of Helsinki. All experiments were performed in compliance with relevant Japanese laws and institutional guidelines. All protocols were approved by the Human Subjects Research Review Committee of the Okinawa Institute of Science and Technology Graduate University (OIST).

### Chemicals and reagents

Standards for metabolite identification were purchased from commercial sources as described previously (Pluskal et al., 2010b) (Chaleckis et al., 2014) (Chaleckis et al., 2016) (Teruya et al., 2019).

### LC-MS analysis and data processing

Urinary metabolites were analyzed using an Ultimate 3000 DGP-3600RS HPLC system (Thermo Fisher Scientific, Waltham, MA, USA) coupled to an LTQ Orbitrap mass spectrometer (Thermo Fisher Scientific, Waltham, MA, USA), as described (Chaleckis et al., 2016) (Kameda et al., 2020). Briefly, LC separation was performed on a ZIC-pHILIC column (Merck SeQuant, Umea, Sweden; 150 mm x 2.1 mm, 5 μm particle size). Acetonitrile (A) and 10 mM ammonium carbonate buffer, pH 9.3 (B) were used as the mobile phase, with a linear gradient elution from 80-20% A over 30 min, at a flow rate of 100 μL mL^-1^. The mass spectrometer was operated in full-scan mode with a 100-1000 m/z scan rate and automatic data-dependent MS/MS fragmentation scans. For each metabolite, we chose a singly charged, [M+H]+ or [M-H]-, peak (**Supplemental Table S3**). Peak detection and identification of metabolites were performed using MZmine 2 software (Pluskal et al., 2010a). Detailed data analytical procedures and parameters have been described previously (Teruya et al., 2019).

### Peak identification and characteristics

We analyzed 99 urine metabolites that were confirmed using standards or MS/MS analysis (Pluskal et al., 2010b) (Chaleckis et al., 2014) (Chaleckis et al., 2016) (Teruya et al., 2019). Metabolites were classified into 3 groups (H, M, and L), according to their peak areas. H denotes compounds with high peak areas (>10^8^ AU), M with medium peak areas (10^7^ ∼10^8^ AU) and L with low peak areas (<10^7^ AU) (**Supplemental Table S2**).

### Statistical analysis

A non-parametric Mann–Whitney test was used to compare young and elderly subjects. Statistical significance was established at p<0.05. Data were exported into a spreadsheet and dot plots were drawn using R statistical software (http://www.r-project.org). Correlation coefficients were determined for identification of metabolic networks among compounds. Principal component analysis (PCA) was conducted using SIMCA-P+ software (Umetrics Inc., Umea, Sweden). Pearson’s correlation coefficients among metabolites were calculated using Microsoft Excel.

## Data availability

Raw LC-MS data in mzML format are accessible via the MetaboLights repository (URL: http://www.ebi.ac.uk/metabolights). Data for the 27 volunteers are available under accession number MTBLS1407.

## Acknowledgments

We thank Ms. Junko Takada for providing excellent technical assistance. We gratefully acknowledge the editorial help of Dr. Steven D. Aird. We are greatly indebted to the generous support of Okinawa Institute of Science and Technology Graduate University and its Innovative Technology Research (ITR) fund.

## Supplementary information

**Supplementary Table S1.**
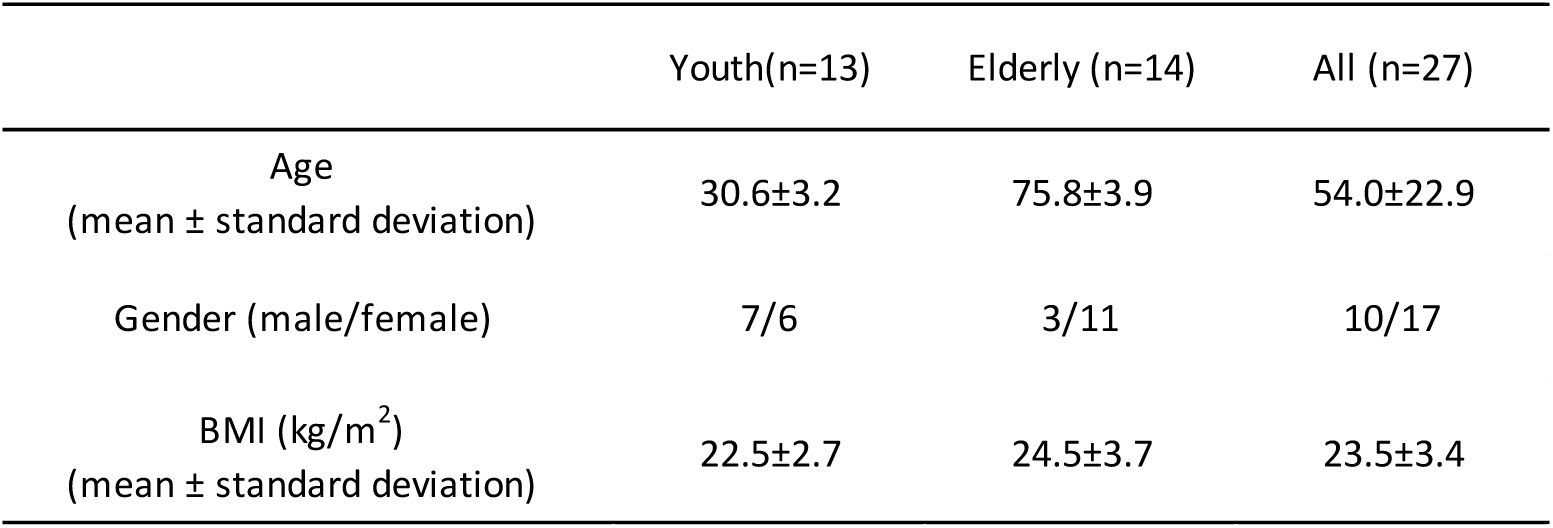
Characteristics of 27 subjects

**Supplemental Table S2.**
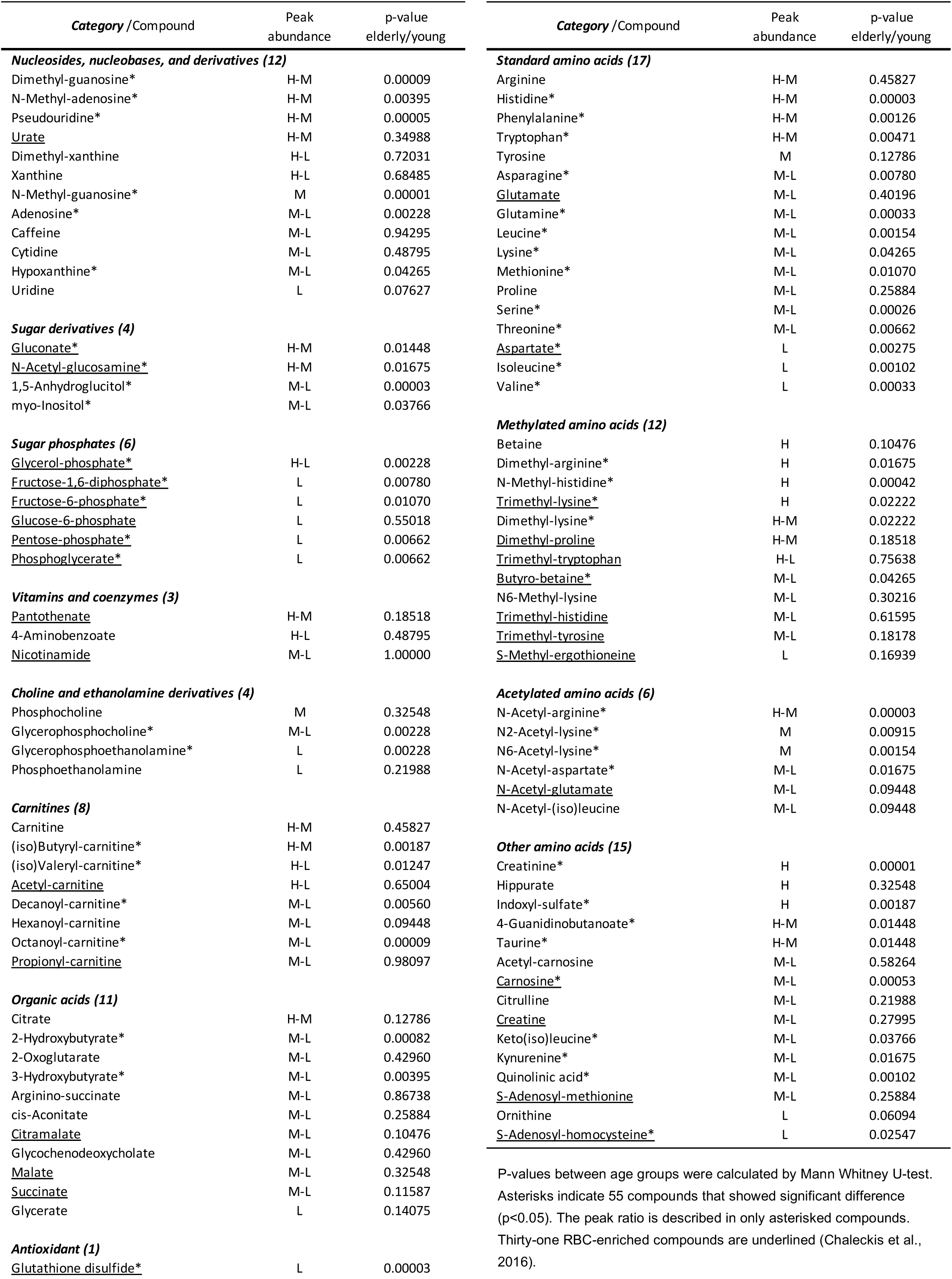
List of 99 urine compounds identified in the present study.

**Supplemental Table S3.**
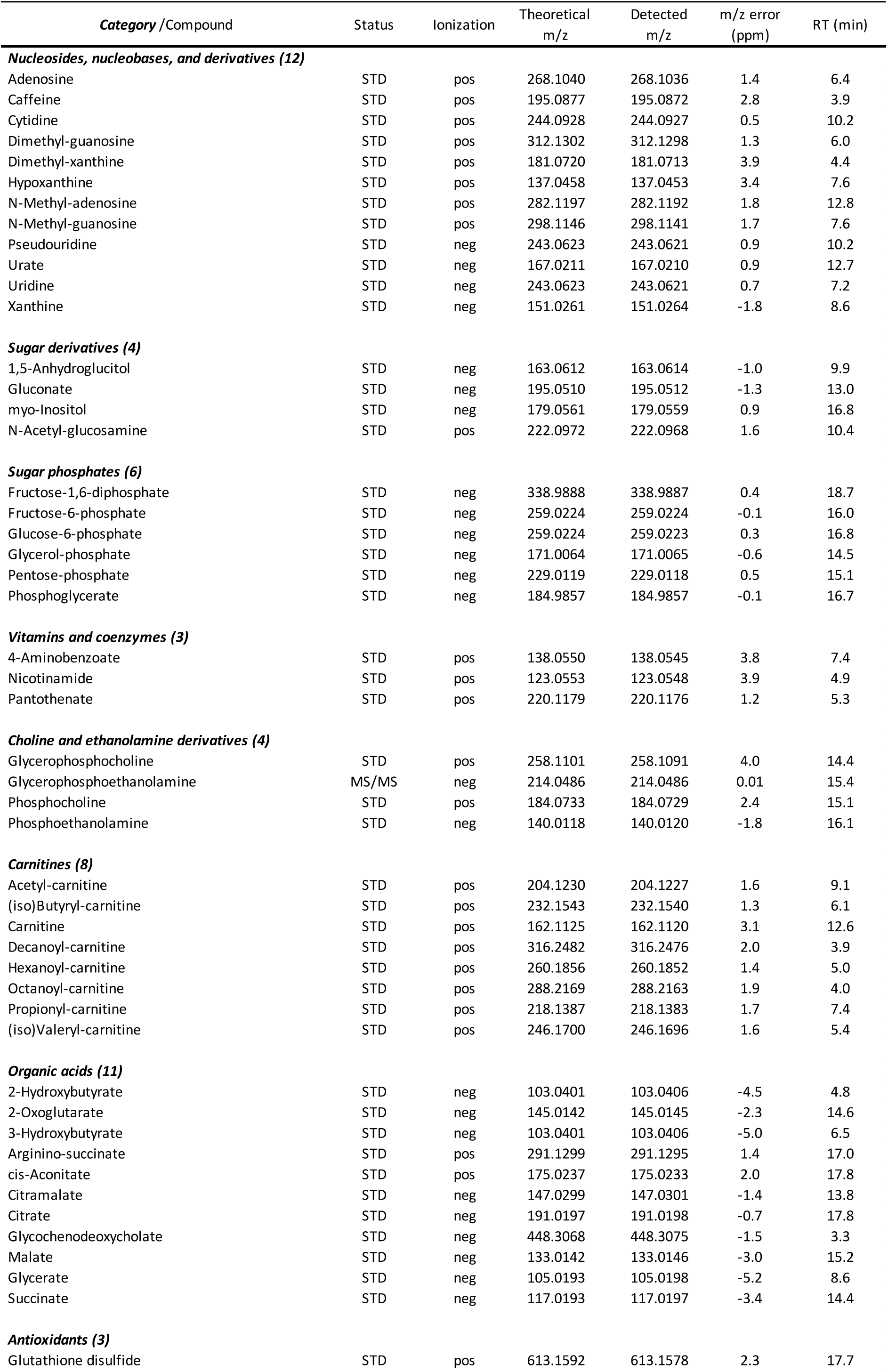
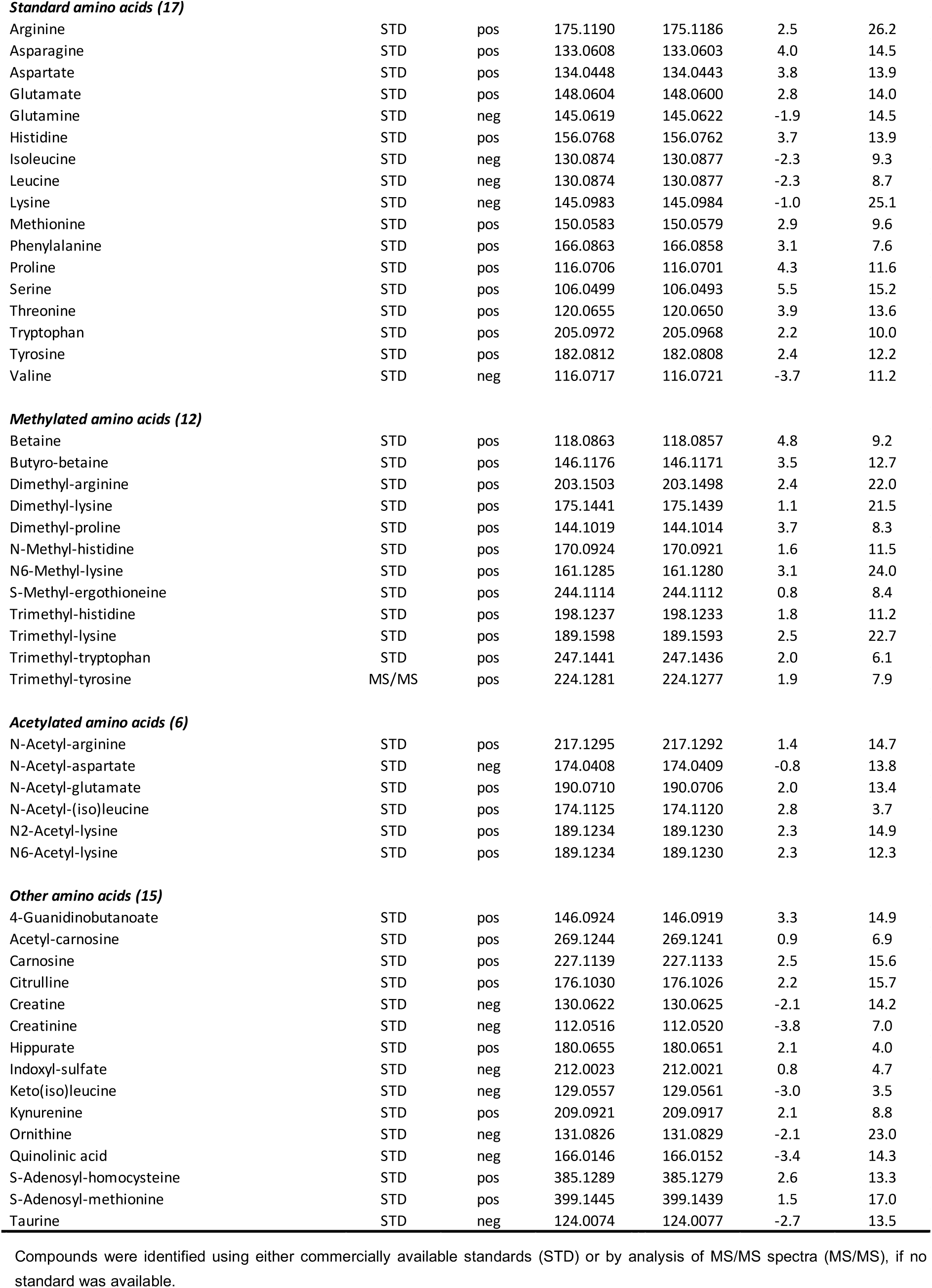
Chromatogram and mass spectrum data.

| *Category* /Compound | Status | Ionization | Theoretical<br>m/z | Detected<br>m/z | m/z error<br>(ppm) | RT (min) |
| --- | --- | --- | --- | --- | --- | --- |
| ***Nucleosides, nucleobases, and derivatives (12)*** |  |  |  |  |  |  |
| Adenosine | STD | pos | 268.1040 | 268.1036 | 1.4 | 6.4 |
| Caffeine | STD | pos | 195.0877 | 195.0872 | 2.8 | 3.9 |
| Cytidine | STD | pos | 244.0928 | 244.0927 | 0.5 | 10.2 |
| Dimethyl-guanosine | STD | pos | 312.1302 | 312.1298 | 1.3 | 6.0 |
| Dimethyl-xanthine | STD | pos | 181.0720 | 181.0713 | 3.9 | 4.4 |
| Hypoxanthine | STD | pos | 137.0458 | 137.0453 | 3.4 | 7.6 |
| N-Methyl-adenosine | STD | pos | 282.1197 | 282.1192 | 1.8 | 12.8 |
| N-Methyl-guanosine | STD | pos | 298.1146 | 298.1141 | 1.7 | 7.6 |
| Pseudouridine | STD | neg | 243.0623 | 243.0621 | 0.9 | 10.2 |
| Urate | STD | neg | 167.0211 | 167.0210 | 0.9 | 12.7 |
| Uridine | STD | neg | 243.0623 | 243.0621 | 0.7 | 7.2 |
| Xanthine | STD | neg | 151.0261 | 151.0264 | -1.8 | 8.6 |
| ***Sugar derivatives (4)*** |  |  |  |  |  |  |
| 1,5-Anhydroglucitol | STD | neg | 163.0612 | 163.0614 | -1.0 | 9.9 |
| Gluconate | STD | neg | 195.0510 | 195.0512 | -1.3 | 13.0 |
| myo-Inositol | STD | neg | 179.0561 | 179.0559 | 0.9 | 16.8 |
| N-Acetyl-glucosamine | STD | pos | 222.0972 | 222.0968 | 1.6 | 10.4 |
| ***Sugar phosphates (6)*** |  |  |  |  |  |  |
| Fructose-1,6-diphosphate | STD | neg | 338.9888 | 338.9887 | 0.4 | 18.7 |
| Fructose-6-phosphate | STD | neg | 259.0224 | 259.0224 | -0.1 | 16.0 |
| Glucose-6-phosphate | STD | neg | 259.0224 | 259.0223 | 0.3 | 16.8 |
| Glycerol-phosphate | STD | neg | 171.0064 | 171.0065 | -0.6 | 14.5 |
| Pentose-phosphate | STD | neg | 229.0119 | 229.0118 | 0.5 | 15.1 |
| Phosphoglycerate | STD | neg | 184.9857 | 184.9857 | -0.1 | 16.7 |
| ***Vitamins and coenzymes (3)*** |  |  |  |  |  |  |
| 4-Aminobenzoate | STD | pos | 138.0550 | 138.0545 | 3.8 | 7.4 |
| Nicotinamide | STD | pos | 123.0553 | 123.0548 | 3.9 | 4.9 |
| Pantothenate | STD | pos | 220.1179 | 220.1176 | 1.2 | 5.3 |
| ***Choline and ethanolamine derivatives (4)*** |  |  |  |  |  |  |
| Glycerophosphocholine | STD | pos | 258.1101 | 258.1091 | 4.0 | 14.4 |
| Glycerophosphoethanolamine | MS/MS | neg | 214.0486 | 214.0486 | 0.01 | 15.4 |
| Phosphocholine | STD | pos | 184.0733 | 184.0729 | 2.4 | 15.1 |
| Phosphoethanolamine | STD | neg | 140.0118 | 140.0120 | -1.8 | 16.1 |
| ***Carnitines (8)*** |  |  |  |  |  |  |
| Acetyl-carnitine | STD | pos | 204.1230 | 204.1227 | 1.6 | 9.1 |
| (iso)Butyryl-carnitine | STD | pos | 232.1543 | 232.1540 | 1.3 | 6.1 |
| Carnitine | STD | pos | 162.1125 | 162.1120 | 3.1 | 12.6 |
| Decanoyl-carnitine | STD | pos | 316.2482 | 316.2476 | 2.0 | 3.9 |
| Hexanoyl-carnitine | STD | pos | 260.1856 | 260.1852 | 1.4 | 5.0 |
| Octanoyl-carnitine | STD | pos | 288.2169 | 288.2163 | 1.9 | 4.0 |
| Propionyl-carnitine | STD | pos | 218.1387 | 218.1383 | 1.7 | 7.4 |
| (iso)Valeryl-carnitine | STD | pos | 246.1700 | 246.1696 | 1.6 | 5.4 |
| ***Organic acids (11)*** |  |  |  |  |  |  |
| 2-Hydroxybutyrate | STD | neg | 103.0401 | 103.0406 | -4.5 | 4.8 |
| 2-Oxoglutarate | STD | neg | 145.0142 | 145.0145 | -2.3 | 14.6 |
| 3-Hydroxybutyrate | STD | neg | 103.0401 | 103.0406 | -5.0 | 6.5 |
| Arginino-succinate | STD | pos | 291.1299 | 291.1295 | 1.4 | 17.0 |
| cis-Aconitate | STD | pos | 175.0237 | 175.0233 | 2.0 | 17.8 |
| Citramalate | STD | neg | 147.0299 | 147.0301 | -1.4 | 13.8 |
| Citrate | STD | neg | 191.0197 | 191.0198 | -0.7 | 17.8 |
| Glycochenodeoxycholate | STD | neg | 448.3068 | 448.3075 | -1.5 | 3.3 |
| Malate | STD | neg | 133.0142 | 133.0146 | -3.0 | 15.2 |
| Glycerate | STD | neg | 105.0193 | 105.0198 | -5.2 | 8.6 |
| Succinate | STD | neg | 117.0193 | 117.0197 | -3.4 | 14.4 |
| ***Antioxidants (3)*** |  |  |  |  |  |  |
| Glutathione disulfide | STD | pos | 613.1592 | 613.1578 | 2.3 | 17.7 |
| ***Standard amino acids (17)*** |  |  |  |  |  |  |
| Arginine | STD | pos | 175.1190 | 175.1186 | 2.5 | 26.2 |
| Asparagine | STD | pos | 133.0608 | 133.0603 | 4.0 | 14.5 |
| Aspartate | STD | pos | 134.0448 | 134.0443 | 3.8 | 13.9 |
| Glutamate | STD | pos | 148.0604 | 148.0600 | 2.8 | 14.0 |
| Glutamine | STD | neg | 145.0619 | 145.0622 | -1.9 | 14.5 |
| Histidine | STD | pos | 156.0768 | 156.0762 | 3.7 | 13.9 |
| Isoleucine | STD | neg | 130.0874 | 130.0877 | -2.3 | 9.3 |
| Leucine | STD | neg | 130.0874 | 130.0877 | -2.3 | 8.7 |
| Lysine | STD | neg | 145.0983 | 145.0984 | -1.0 | 25.1 |
| Methionine | STD | pos | 150.0583 | 150.0579 | 2.9 | 9.6 |
| Phenylalanine | STD | pos | 166.0863 | 166.0858 | 3.1 | 7.6 |
| Proline | STD | pos | 116.0706 | 116.0701 | 4.3 | 11.6 |
| Serine | STD | pos | 106.0499 | 106.0493 | 5.5 | 15.2 |
| Threonine | STD | pos | 120.0655 | 120.0650 | 3.9 | 13.6 |
| Tryptophan | STD | pos | 205.0972 | 205.0968 | 2.2 | 10.0 |
| Tyrosine | STD | pos | 182.0812 | 182.0808 | 2.4 | 12.2 |
| Valine | STD | neg | 116.0717 | 116.0721 | -3.7 | 11.2 |
| ***Methylated amino acids (12)*** |  |  |  |  |  |  |
| Betaine | STD | pos | 118.0863 | 118.0857 | 4.8 | 9.2 |
| Butyro-betaine | STD | pos | 146.1176 | 146.1171 | 3.5 | 12.7 |
| Dimethyl-arginine | STD | pos | 203.1503 | 203.1498 | 2.4 | 22.0 |
| Dimethyl-lysine | STD | pos | 175.1441 | 175.1439 | 1.1 | 21.5 |
| Dimethyl-proline | STD | pos | 144.1019 | 144.1014 | 3.7 | 8.3 |
| N-Methyl-histidine | STD | pos | 170.0924 | 170.0921 | 1.6 | 11.5 |
| N6-Methyl-lysine | STD | pos | 161.1285 | 161.1280 | 3.1 | 24.0 |
| S-Methyl-ergothioneine | STD | pos | 244.1114 | 244.1112 | 0.8 | 8.4 |
| Trimethyl-histidine | STD | pos | 198.1237 | 198.1233 | 1.8 | 11.2 |
| Trimethyl-lysine | STD | pos | 189.1598 | 189.1593 | 2.5 | 22.7 |
| Trimethyl-tryptophan | STD | pos | 247.1441 | 247.1436 | 2.0 | 6.1 |
| Trimethyl-tyrosine | MS/MS | pos | 224.1281 | 224.1277 | 1.9 | 7.9 |
| ***Acetylated amino acids (6)*** |  |  |  |  |  |  |
| N-Acetyl-arginine | STD | pos | 217.1295 | 217.1292 | 1.4 | 14.7 |
| N-Acetyl-aspartate | STD | neg | 174.0408 | 174.0409 | -0.8 | 13.8 |
| N-Acetyl-glutamate | STD | pos | 190.0710 | 190.0706 | 2.0 | 13.4 |
| N-Acetyl-(iso)leucine | STD | pos | 174.1125 | 174.1120 | 2.8 | 3.7 |
| N2-Acetyl-lysine | STD | pos | 189.1234 | 189.1230 | 2.3 | 14.9 |
| N6-Acetyl-lysine | STD | pos | 189.1234 | 189.1230 | 2.3 | 12.3 |
| ***Other amino acids (15)*** |  |  |  |  |  |  |
| 4-Guanidinobutanoate | STD | pos | 146.0924 | 146.0919 | 3.3 | 14.9 |
| Acetyl-carnosine | STD | pos | 269.1244 | 269.1241 | 0.9 | 6.9 |
| Carnosine | STD | pos | 227.1139 | 227.1133 | 2.5 | 15.6 |
| Citrulline | STD | pos | 176.1030 | 176.1026 | 2.2 | 15.7 |
| Creatine | STD | neg | 130.0622 | 130.0625 | -2.1 | 14.2 |
| Creatinine | STD | neg | 112.0516 | 112.0520 | -3.8 | 7.0 |
| Hippurate | STD | pos | 180.0655 | 180.0651 | 2.1 | 4.0 |
| Indoxyl-sulfate | STD | neg | 212.0023 | 212.0021 | 0.8 | 4.7 |
| Keto(iso)leucine | STD | neg | 129.0557 | 129.0561 | -3.0 | 3.5 |
| Kynurenine | STD | pos | 209.0921 | 209.0917 | 2.1 | 8.8 |
| Ornithine | STD | neg | 131.0826 | 131.0829 | -2.1 | 23.0 |
| Quinolinic acid | STD | neg | 166.0146 | 166.0152 | -3.4 | 14.3 |
| S-Adenosyl-homocysteine | STD | pos | 385.1289 | 385.1279 | 2.6 | 13.3 |
| S-Adenosyl-methionine | STD | pos | 399.1445 | 399.1439 | 1.5 | 17.0 |
| Taurine | STD | neg | 124.0074 | 124.0077 | -2.7 | 13.5 |

Compounds were identified using either commercially available standards (STD) or by analysis of MS/MS spectra (MS/MS), if no standard was available.

**Supplemental Figure S1.**
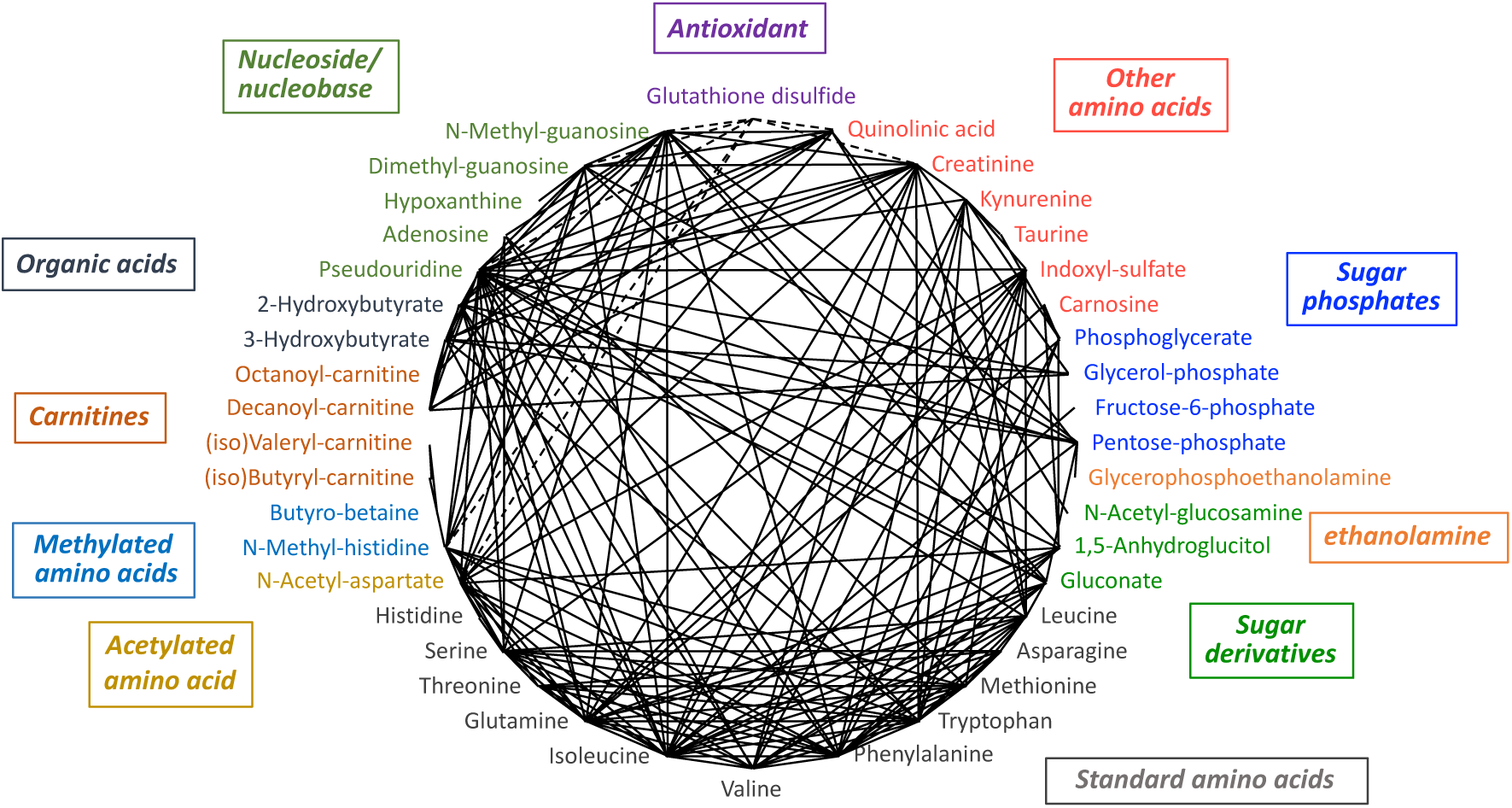
Of the 55 aging markers, 13 compounds with correlations <0.7. The 5 most highly correlated compounds for each.

**Supplemental Figure S2.**
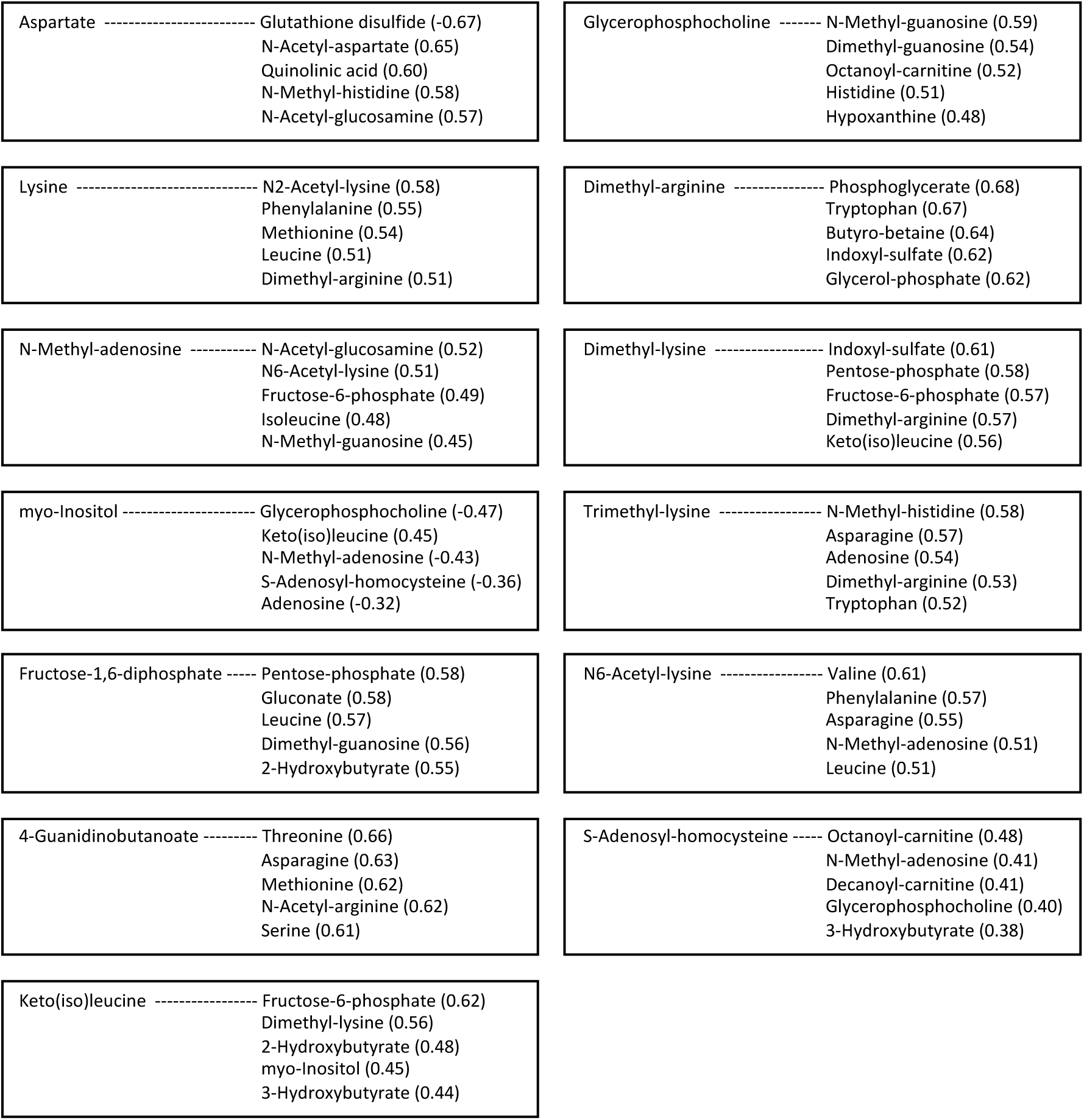
Of the 55 aging markers, 13 compounds with correlations <0.7. The 5 most highly correlated compounds for each.

